# The role of extracellular matrix biomolecules on endometrial epithelial cell attachment and cytokeratin 18 expression on gelatin hydrogels

**DOI:** 10.1101/2021.10.24.465574

**Authors:** Samantha G. Zambuto, Ishita Jain, Kathryn B.H. Clancy, Gregory H. Underhill, Brendan A.C. Harley

## Abstract

The endometrium undergoes profound changes in tissue architecture and composition, both during the menstrual cycle as well as in the context of pregnancy. Dynamic remodeling processes of the endometrial extracellular matrix (ECM) are a major element of endometrial homeostasis, including changes across the menstrual cycle. A critical element of this tissue microenvironment is the endometrial basement membrane, a specialized layer of proteins that separates the endometrial epithelium from the underlying endometrial ECM. Bioengineering models of the endometrial microenvironment that present an appropriate endometrial ECM and basement membrane may provide an improved environment to study endometrial epithelial cell (EEC) function. Here, we exploit a tiered approach using two-dimensional high throughput microarrays and three-dimensional gelatin hydrogels to define patterns of EEC attachment and cytokeratin 18 (CK18) expression in response to combinations of endometrial basement membrane proteins. We identify combinations (collagen IV + tenascin C; collagen I + collagen III; hyaluronic acid + tenascin C; collagen V; collagen V + hyaluronic acid; collagen III; collagen I) that facilitate increased EEC attachment, increased CK18 intensity, or both. We also identify significant EEC mediated remodeling of the GelMA matrix environment via analysis of nascent protein deposition. Together, we report efforts to tailor the localization of basement membrane-associated proteins and proteoglycans in order to investigate tissue engineered models of the endometrial microenvironment.

## 1. Introduction

The endometrium is the lining of the uterus [1]. The endometrium is stratified, with an epithelium overlaying a highly vascularized stromal compartment [1]. The luminal epithelium is the location of critical cell-cell dialogue between endometrial cells and an implanting blastocyst [1]. The luminal epithelium also remodels during each menstrual cycle phase to become more receptive to a potential pregnancy [1]. A basement membrane layer connects the epithelial and stromal layers; although an expanding group of extracellular matrix and adhesion molecules have been identified as part of the basement membrane, much remains unknown regarding the dynamic remodeling processes that occur at its intersection with the endometrial epithelium [1, 2]. The study of many of these processes is intractable *in vivo* due to our inability to study early pregnancy in humans and due to significant differences between pregnancy in humans and the most common animal models [9–11]. *In vitro* endometrial model systems provide a framework to investigate physiological processes underlying endometrial activity. Two-dimensional models of the endometrial epithelium fail to recapitulate the native 3D tissue structure. Further, many existing endometrial epithelial models do not incorporate a basement membrane layer despite its importance in endometrial function and pregnancy [3–5]. A critical opportunity exists to develop engineered systems to examine the role of ECM biomolecules on the dynamic nature of the endometrial epithelium and its basement membrane. Such a platform would provide an important framework to investigate the role of the basement membrane on endometrial epithelial cell attachment and phenotype.

First generation models of the endometrial basement microenvironment have included a range of conventional hydrogel models, notably Matrigel™, fibrin, alginate, and hyaluronic gels. Matrigel™, a commonly used ECM mixture used to mimic the basement membrane, consists of a mixture of thousands of proteins, primarily type IV collagen, laminin, and nidogen, derived from Engelbrecht-Holm Swarm (EHS) murine sarcoma cells [8]. However, lot variability and a heterogeneous protein composition, along with an inability to decouple biophysical and biochemical properties, render it difficult to examine matrix-templated endometrial cell activity [8, 12, 13]. Despite the diversity of proteins explicitly linked to the endometrium, the majority of prior studies use generic basement membrane constituents such as fibronectin or collagen IV and laminin [3] and have not explored a wider range of combinations. As a result, we propose a systematic examination of combinations of known biomolecules within the endometrium to determine the roles they may play on epithelial cellular behavior.

We previously described a first generation stratified endometrial culture formed via seeding of primary endometrial epithelial cell (EEC) cultures on methacrylamide-functionalized gelatin (GelMA) hydrogels [14]. Because gelatin contains RGD cell binding motifs [15], EECs attached to the GelMA hydrogel surface without any additional basement membrane layer; however, we observed that cells formed non-uniform layers on the gels that lost stability over time. Strategies to immobilize supplemental basement membrane proteins onto defined endometrial hydrogel formulations may allow for the inclusion of defined basement membrane layer analogs the provide the ability to locally control ligand density and protein/proteoglycan combinations to more faithfully replicate endometrial basement membrane function. Recent advances with microbial transglutaminase (mTg) suggest a route to enzymatically crosslink a layer of ECM proteins onto a gel surface [16], offering a strategy to decouple endometrial basement membrane properties from those of the underlying endometrial ECM-inspired hydrogel environment. Furthermore, mTg is water-soluble, inexpensive, biocompatible, and stable long term which renders it an attractive method for producing coated GelMA hydrogels for long term cell culture [16].

The endometrial extracellular matrix exhibits dynamic changes in tissue composition and architecture during homeostasis, pregnancy, and in response to endometrial-associated pathologies [1]. Previous studies have identified a wide range of ECM-associated proteins and proteoglycans in the endometrium and decidua whose expression changes markedly across the menstrual cycle or in response to steroidal sex hormones [1]. For our study, we systematically investigated the influence of combinations of 10 extracellular matrix proteins and proteoglycans present in the endometrium on endometrial epithelial cell activity: collagens I, III, IV, V, decorin, fibronectin, hyaluronic acid, laminin, lumican, tenascin C. These biomolecules were chosen because of their functional significance in structural remodeling, tissue biomechanics, cell adhesion, cell proliferation, tissue differentiation, as well as other cellular processes crucial to endometrial remodeling during the menstrual cycle and pregnancy [1]. Collagen I is most prevalent in the endometrium throughout the menstrual cycle as well as during pregnancy [1, 6] while collagen III is present during all menstrual cycle phases but less abundant than collagen I during the first trimester of pregnancy [1, 6]. Collagen IV is present during the menstrual cycle [6] while collagen V increases during decidualization [1]. Fibronectin is highly abundant in the endometrium and decidual ECM [1] while tenascin C is more localized near stromal cells surrounding proliferating or developing endometrium epithelia [1] and decorin is present in decidual ECM [1]. Although primarily studied in the context of murine models, lumican is localized within the uterine stroma during pregnancy and may potentially be important in human pregnancy as well [1]. Laminin is present in the decidua and mediates trophoblast attachment and spreading [1] while hyaluronic acid likely influences elevated hydration during the mid-proliferative and mid-secretory phases in the human endometrium [1]. The basement membrane plays an important role in mucosal barrier tissues, particularly the endometrium [1, 7, 8]. In humans, the decidual basement membrane consists of laminin, collagens type IV, VII, XVII, and heparan sulfate proteoglycans [1]. Because of the dynamic nature of the endometrial ECM and basement membrane, tissue engineering models offer a unique opportunity to examine the role of endometrial ECM and basement membrane factors in endometrial cell activity, remodeling, and processes associated with trophoblast invasion and placentation.

We report a tissue engineering approach to examine benchmarks of endometrial epithelial cell activity (e.g., cell attachment, phenotypic markers of attachment). We describe a high throughput microarray-based approach to investigate shifts in EEC attachment and epithelial maturation in response to 55 single and pairwise combinations of 10 ECM proteins and proteoglycans found in the endometrium ECM (collagens I, III, IV, V, decorin, fibronectin, hyaluronic acid, laminin, lumican, tenascin C). We track the degree of cell attachment as well as a biomarker of epithelial monolayer maturation (expression of cytokeratin 18, CK18) responsible for anchoring the endometrial epithelial cell cytoskeleton to the basement membrane [3]. We identified ECM combinations that promote both high and moderate levels of cell attachment and CK18 intensity, combinations that promoted high cell attachment and median CK18 intensity, and combinations that promote median cell attachment and high CK18 intensity. We subsequently used microbial transglutaminase to immobilize a down-selected set of ECM combinations on three-dimensional methacrylamide-functionalized gelatin (GelMA) hydrogels. We subsequently evaluate patterns of attachment, CK18 expression, and nascent protein deposition by EECs on GelMA hydrogels. This project describes a pipeline for creating and characterizing model basement membrane constructs as part of a larger tissue engineered strategy to replicate the stratified endometrium.

## 2. Materials and Methods

### 2.1 Cell Culture and Maintenance

We cultured primary human endometrial epithelial cells (EECs; LifeLine Cell Technology FC-0078) as per the manufacturer’s instructions and used them experimentally at two passages from receipt. Cells were cultured in phenol red-free medium (LifeLine Cell Technology) and in 5% CO_2_ incubators at 37°C. Cells were routinely tested for mycoplasma contamination using the MycoAlert™ Mycoplasma Detection Kit (Lonza).

### 2.2. Microarray Fabrication and Experimentation

#### 2.2.1. Microarray Fabrication

We prepared polyacrylamide (PA) hydrogels following previous protocols [17–19]. Briefly, 25×75 mm glass microscope slides were washed with 0.25% v/v Triton X-100 in dH_2_O and placed on an orbital shaker for 30 minutes. After rinsing with dH_2_O, slides were immersed in acetone and methanol for 30 minutes each on an orbital shaker. This was followed by etching the glass slide by immersing them 0.2 N NaOH for 1 hour on an orbital shaker and then rinsing with dH_2_O. The slides were then air-dried and placed on a hot plate at 110°C until dry. For silanization, the cleaned slides were immersed in 2% v/v 3-(trimethoxysilyl)propyl methacrylate in ethanol and placed on the shaker for 30 minutes, followed by a wash in ethanol for 5 minutes. The silanized slides were air-dried, and again placed on the hot plate at 110°C until dry. For fabrication of hydrogels with specific elastic moduli, prepolymer solution with 6% acrylamide (Sigma-Aldrich A3553-100G) and 0.45% bis-acrylamide (Sigma-Aldrich M7279-25G) was prepared to achieve elastic moduli of 6 kPa. The prepolymer solution was then mixed with Irgacure 2959 (BASF, Corp.) solution (20% w/v in methanol) at a final volumetric ratio of 9:1 (prepolymer:Irgacure). This working solution was then deposited onto slides (100 μL/slide) and covered with 22×60 mm cover glasses. The sandwiched working solution was transferred to a UV oven and exposed to 365 nm UV A for 10 min (240E3 μJ). After removing the cover glasses, the slides were immersed in dH_2_O at room temperature for 3 days in order to remove excess reagents from the hydrogel substrates. Before microarray fabrication, hydrogel substrates were thoroughly dehydrated on a hot plate for ≥15 minutes at 50°C. Microarrays were fabricated as described previously [20–22]. Extracellular matrix proteins for arraying were diluted in 2×growth factor buffer (38% v/v glycerol in 1× phosphate-buffered saline (PBS), 10.55 mg/mL sodium acetate, 3.72 mg/mL EDTA, 10 mg/mL CHAPS) to a final concentration of 250 μg/mL and loaded in a 384-well V-bottom microplate. List of all the ECM venders and part numbers can be found in Table 1. A robotic benchtop microarrayer (OmniGrid Micro, Digilab) loaded with SMPC Stealth microarray pins (ArrayIt) was used to microprint ECM combinations from the 384 microplate to polyacrylamide hydrogel substrate, resulting in ~600 μm diameter arrayed domains. Fabricated arrays were stored at room temperature and 65% RH overnight and left to dry under ambient conditions in the dark.

**Table 1.** Microarray Biomolecule Information.

| **Biomolecule** | **Vendor** | **Catalog Number** | **Stock Concentration** |
| --- | --- | --- | --- |
| Collagen 1 | EMD Millipore | 08-115MI | 1 mg/mL |
| Collagen 3 | EMD Millipore | CC054 | 1 mg/mL |
| Collagen 4 | Abcam | ab7536 | 1 mg/mL |
| Collagen 5 | Abcam | ab7530 | 1 mg/mL |
| Fibronectin | EMD Millipore | FC010-10MG | 1 mg/mL |
| Decorin | R&D Systems | 143-DE-100 | 0.5 mg/mL |
| Lumican | ACROBiosystems | LUM-H5227-100ug | 0.5 mg/mL |
| Laminin | EMD Millipore | CC095 | 1 mg/mL |
| Hyaluronic Acid | Lifecore Biomedical | HA60k-1 | 1 mg/mL |
| Tenascin C | R&D Systems | 3358-TC-050 | 0.5 mg/mL |

#### 2.2.2. Microarray Cell Culture

We sterilized microarrays using a 1% penicillin/streptomycin (Thermo Fisher) solution for 20 minutes under UV light. EECs were then passaged and seeded at a density of 500,000 cells per microarray in 4 mL cell growth medium. After seeding, microarrays were shaken by hand every 30 minutes for 2 hours and then rinsed with cell growth medium 6 hours after seeding. Microarrays were cultured in slide plates for 24 hours after seeding and cultured in 5% CO_2_ incubators at 37°C. Three independent biological replicates were cultured and analyzed for these experiments with at least three technical replicates per experiment.

#### 2.2.3. Microarray Staining

We fixed microarray cultures 24 hours after seeding using 4 mL of formalin (Sigma-Aldrich) followed by three PBS washes. Cultures were stored in PBS at 4°C until use. Microarrays were permeabilized in a 0.5% Tween20 (Fisher Scientific) solution for 15 minutes followed by three 5 minute washes in (PBST). Samples were blocked for 1 hour at room temperature in a 2% Abdil solution (bovine serum albumin; Sigma Aldrich, Tween20, PBS) and subsequently incubated in primary antibody (cytokeratin 18; 1:250; Abcam ab52948) overnight at 4°C. Samples were then washed three times with PBST and then cultured in secondary antibody (Alexafluor 488 goat anti-rabbit; 1:500; Thermo Fisher A-11008) overnight at 4°C. Three 5 minute PBST washes were performed and then slides were mounted with DAPI Fluoromount (Southern Biotechnology Associates, Inc.).

#### 2.3.4. Microarray Imaging

We imaged the microarrays using Axioscan.Z1 Slide Scanner and 10X objective. A wide tile region was defined for the whole array region which was then stitched offline using Zen and exported into TIFF Images for each individual channel.

#### 2.2.5. Microarray Image Analysis

We converted images of entire arrays to individual 8-bit TIFF files per channel (i.e., red, green, blue, and gray) by Fiji (ImageJ version 1.52p) [23]. CellProfiler (version 4.0.0) was used obtain per cell measurement for each channel [24]. The images were cropped in MATLAB (version R2018b) to separate each array in a single image. Positional information for each array was automatically calculated using their relative position from the positional dextran-rhodamine markers. Nuclei were identified using the DAPI channel image using IdentifyPrimaryObject module and cell boundary was identified using the CK18 stain around these nuclei using IdentifySecondaryObject module. The MeasureObjectIntensity module was used to quantify single-cell intensity. The data were exported to CSV files that were then imported in RStudio for data visualization.

### 2.3. Methacrylamide-Functionalized Gelatin (GelMA) Hydrogel Experimentation

#### 2.3.1. Synthesis and Fabrication of GelMA Hydrogels

GelMA was synthesized as described previously and was found to have a degree of functionalization of 57%, determined via ^1^H-NMR [15, 25]. Prior to hydrogel fabrication, lyophilized GelMA was sterilized under UV light for 30 minutes. To fabricate hydrogels, a solution was created consisting of lyophilized GelMA (5 wt%) combined with 1% fluorescent beads (Fluospheres Polystyrene Microspheres 1.0 μm orange fluorescent beads; 1×10^10^ beads/mL; Invitrogen) and dissolved at 37°C in phosphate buffered saline (PBS; Lonza 17-516F). Microspheres were used to visualize hydrogels during imaging. 0.1% w/v lithium acylphosphinate (LAP) was added to this solution as a photoinitiator. 10 μL of the prepolymer solution was added to each well of Ibidi μ-Slides Angiogenesis Glass Bottom and was polymerized under UV light (λ=365 nm, 7.14 mW cm^−2^; AccuCure Spot System ULM-3-365) for 30 s.

#### 2.3.2. Extracellular Matrix Coating of GelMA Hydrogels

We used microbial transglutaminase (mTg; Zedira T001, Lot 0920aT001) to coat polymerized GelMA hydrogels with extracellular matrix (ECM) proteins [16, 26, 27]. A 1:1 ratio of 0.5 mg/mL mTg and 10 μg/mL ECM protein were combined and 20 μL of this solution was pipetted onto hydrogels. If two ECM proteins were used to coat gels, they were combined in a 1:1 ratio. Coated hydrogels were incubated for 1 hour in 5% CO_2_ incubators at 37°C. A quick wash was performed using 20 μL of PBS. Information regarding ECM protein vendors and part numbers can be found in Table 1.

#### 2.3.3. Visualization of ECM Coated Hydrogels

We evaluated the effectiveness of mTg-based matrix immobilization for Laminin-coated GelMA hydrogels. Immediately following laminin coating, hydrogels were blocked for 1 hour in 30 μL of a 2% Abdil solution (bovine serum albumin, Tween20, PBS). Samples were then stained with anti-laminin primary antibody (1:200, 30 μL; Abcam ab11575) at 4°C overnight. Three PBS washes were performed followed by incubation with secondary antibody (1:500, 30 μL; Alexafluor 488 goat anti-rabbit Thermo Fisher A-11008) for 2 hours at room temperature. Three additional PBS washes were performed and samples were stored in PBS at 4°C until imaged. Samples were imaged using a DMi8 Yokogawa W1 spinning disc confocal microscope outfitted with a Hamamatsu EM-CCD digital camera (Leica Microsystems). Two fluorescent Z-stacks were taken per gel using the 10x objective, 10 μm step size, and 50-100 slices (3 hydrogels per condition). Maximum intensity image projections were created using FIJI.

### 2.4. Endometrial Epithelial Cell Hydrogel Cultures

#### 2.4.1. Endometrial Epithelial Cell Hydrogels

We fabricated and coated hydrogels as described above, with EECs subsequently seeded at 200,000 cells/cm^2^ onto the hydrogel surface. EECs were allowed to adhere to the hydrogels for 1 hour and were subsequently washed with 50 μL cell medium to remove unattached cells. Hydrogels were cultured for 3 days in 5% CO2 incubators at 37°C and culture medium was replaced daily.

#### 2.4.3. Immunofluorescent Staining

Hydrogels were fixed on day 3 of culture using 4 mL of formalin (Sigma-Aldrich) followed by three PBS washes. Hydrogels were permeabilized in a 0.5% Tween20 (Fisher Scientific) solution for 15 minutes followed by three 5 minute washes in (PBST). Samples were blocked for 1 hour at room temperature in a 2% Abdil solution (bovine serum albumin; Sigma Aldrich, Tween20, PBS) and subsequently incubated in primary antibody (cytokeratin 18; 1:250; Abcam ab52948) overnight at 4°C. Hydrogels were washed 4 × 20 minutes with PBST and then cultured in secondary antibody (Alexafluor 488 goat anti-rabbit; 1:500; Thermo Fisher A-11008) overnight at 4°C. Hydrogels were washed 4 × 20 minutes with PBST and then incubated in Phalloidin-iFluor 633 Reagent (Abcam ab176758) as per the manufacturer’s instructions for 90 minutes at room temperature. Cells were rinsed in PBS 4 × 20 minutes and then incubated in Hoechst (1:2000; Thermo Fisher) for 30 minutes. One PBS wash was performed and samples were stored in PBS until imaged.

#### 2.4.4. Hydrogel Confocal Imaging

We imaged hydrogels using a Zeiss LSM 710 Confocal Microscope and 10X objective. One image was taken per hydrogel (n=3 hydrogels) in a random region of interest. Maximum intensity image projections used for analysis were generated using ZEN (black edition; Zeiss).

#### 2.4.5. Image Analysis

We used CellProfiler (version 4.0.0) to analyze CK18 intensity from maximum intensity projection images of cells seeded onto hydrogels [24]. Nuclei were identified using the DAPI channel image using IdentifyPrimaryObject module and cell boundary was identified using the CK18 stain around these nuclei using IdentifySecondaryObject module. The MeasureObjectIntensity module was used to quantify single-cell intensity. The data were exported to CSV files that were then imported in RStudio for data visualization.

### 2.5. Nascent Protein Deposition

#### 2.5.1. Nascent Protein Staining

Following the protocols developed by Loebel et al. [28, 29], we performed metabolic labeling to visualize nascent protein deposition. Briefly, epithelial hydrogel cultures were cultured as described above. On day 3 of culture, hydrogels were washed once with Hanks’ Balanced Salt Solution (HBSS) and incubated in HBSS for 30 minutes to deplete the cells of any remaining methionine. The HBSS was removed and cells were then incubated at 37°C in HBSS containing the methionine analog azidohomoalanine (Click-iT AHA; Invitrogen, C10102; 100 μM) for 1 hour. Following incubation, two HBSS washes were performed and hydrogels were then incubated in AFDye 488 DBCO (Click Chemistry Tools, 1278-1; 30 μM) for 40 minutes at 37°C, washed three times with PBS, fixed for 15 minutes followed by three PBS washes, and stored at 4°C in PBS until staining. Hydrogels were incubated in CellMask™ Deep Red Plasma Membrane Stain (1:1000; Invitrogen, C10046) for 40 minutes at 37°C followed by three PBS washes. Samples were then incubated in Hoechst at room temperature for 30 minutes (1:2000) followed by one PBS wash. Samples were stored at 4°C in PBS until imaged.

#### 2.5.2. Nascent Protein Intensity Quantification

We used CellProfiler to analyze nascent protein intensity from maximum intensity projection images. Hydrogels were imaged using a Zeiss LSM 710 Confocal Microscope and 10X objective. One image was taken per hydrogel (n=3 hydrogels) in a random region of interest. Maximum intensity image projections used for analysis were generated using ZEN (black edition; Zeiss). These images were uploaded to CellProfiler and analyzed as described above.

### 2.6. Statistics

We used RStudio for our statistical analyses. Normality was determined using the Shapiro-Wilkes test and homoscedasticity was determined via Bartlett’s test for normal data or Levene’s test for non-normal data. Data were analyzed using a one-way analysis of variance (ANOVA) and Tukey post hoc test (normal, homoscedastic), Welch’s ANOVA and Games-Howell post hoc test (normal, heteroscedastic), Kruskall-Wallis ANOVA and Dunn’s post hoc test (non-normal, homoscedastic), or Welch’s Heteroscedastic F Test with Trimmed Means and Winsorized Variances and Games-Howell post hoc test (non-normal, heteroscedastic). Significance was defined as *p*<0.05 and data are presented as box plots unless otherwise described.

## 3. Results

### 3.1. High throughput microarray analysis reveals differential epithelial cell attachment and CK18 expression in response to endometrial ECM biomolecule combinations

Epithelial cell attachment and CK18 expression was quantified in response to 55 single and pairwise combinations of 10 ECM proteins and proteoglycans microarrayed onto slides (Fig. 1A). EECs were analyzed 24 hours after seeding onto arrays to assess cell behavior at early culture timepoints. From this experiment, we observed differential attachment and CK18 expression in EECs based on the ECM proteins on which the cells were cultured. For subsequent analysis, we chose combinations of ECM biomolecules with the highest number of cells attached (Fig. 1B,C; Collagen IV + Tenascin C; Collagen I + Collagen III), or the highest value for CK18 intensity (Fig. 1D,E; Hyaluronic Acid + Tenascin C; Collagen V). We also selected combinations with median (~15^th^ position out of 55 rank ordered samples) levels of cell attachment and CK18 intensity (Fig. 1B-F; Collagen I + Hyaluronic Acid; Collagen V + Hyaluronic Acid) as well as combinations that displayed high cell attachment but median CK18 intensity (Fig. 1B-F; Collagen 1) or high CK18 intensity but median cell attachment (Fig. 1B-F; Collagen III).

**Figure 1.**
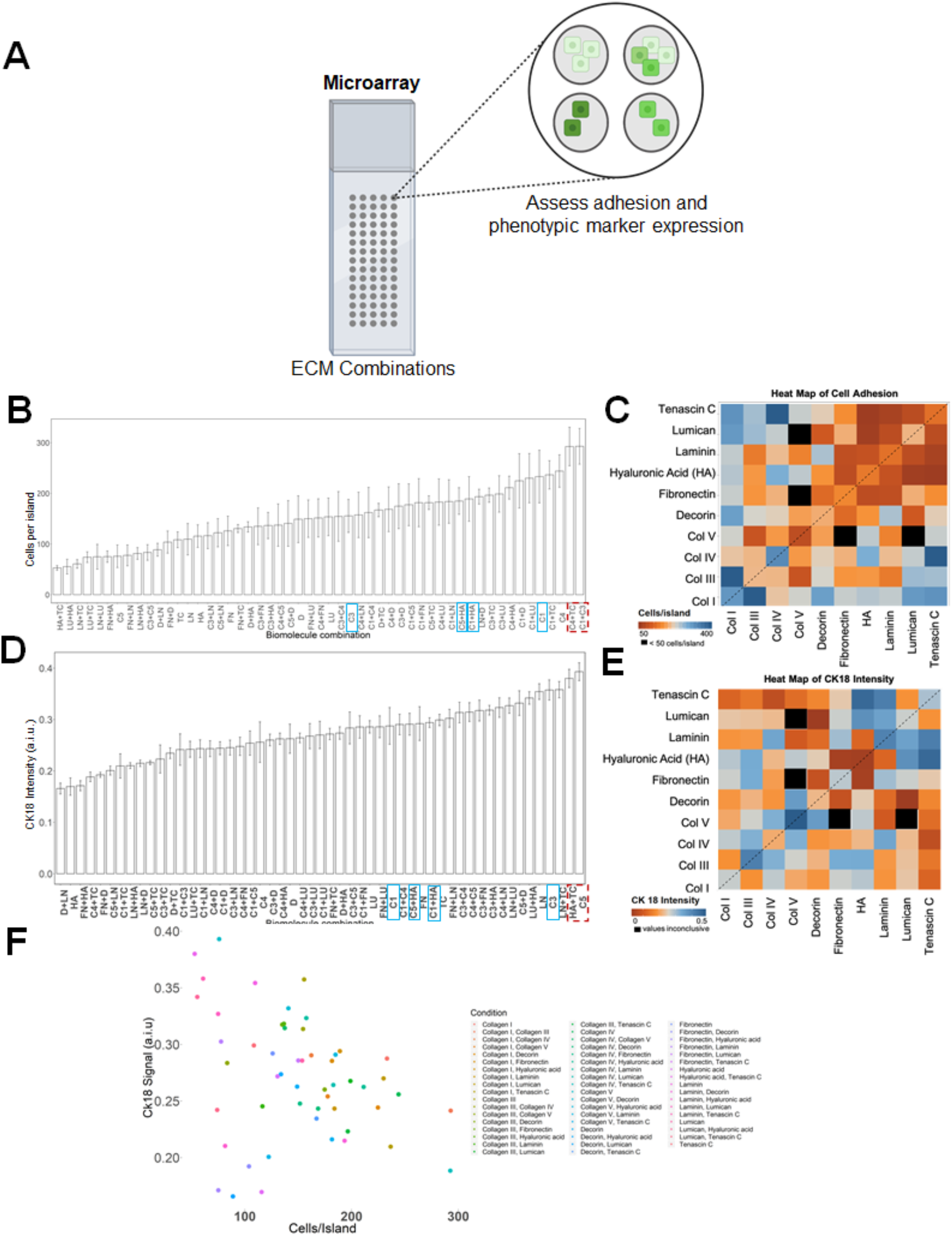
High throughout microarrays demonstrate differences in adhesion and cytokeratin 18 (CK18) intensity. Single and pairwise combinations of extracellular matrix (ECM) biomolecules were arrayed onto polyacrylamide gels to determine adhesion patterns of primary endometrial epithelial cells. Epithelial cells were seeded at a density of 500,000 cells per microarray and cultured for 24 hours prior to fixation and staining. **(A)** Experimental summary. Created with BioRender.com. **(B)** Average number of cells per island on various ECM combinations. Red, dotted box: highest cells per island. Blue boxes: median values of CK18 and adhesion. Data expressed as average ± standard deviation. **(C)** Heat map of cell adhesion based on ECM combinations. **(D)** Average CK18 intensity on various ECM combinations. Red, dotted box: highest CK18 intensity. Blue boxes: median values of CK18 and adhesion. Data expressed as average ± standard deviation. **(E)** Heat map of CK18 intensity based on ECM combinations. **(F)** Scatterplot of average cell adhesion values vs. average CK18 intensity based on ECM combinations. **Key:** C1: Collagen 1; C2: Collagen 2; C3: Collagen 3; C4: Collagen 4; C5: Collagen 5; D: Decorin; FN: Fibronectin; HA: Hyaluronic Acid; LN: Laminin; LU: Lumican; TC: Tenascin C.

### 3.2. GelMA hydrogels can be coated with ECM biomolecules using microbial transglutaminase

Microbial transglutaminase (mTg) was used to coat GelMA hydrogels with proteins/proteoglycan combinations identified via high throughput screen to subsequently investigate EEC attachment and CK18 expression on GelMA hydrogels with functionalized basement membrane layers (Fig. 2A). To confirm biomolecule attachment, GelMA hydrogels were functionalized with laminin, a commonly used ECM protein for cell attachment. Immunofluorescent staining showed that while simple addition of laminin to GelMA hydrogels without mTG resulted in limited laminin adhesion, laminin adhered to GelMA hydrogels via mTg produced a strong laminin signal (Fig. 2B).

**Figure 2.**
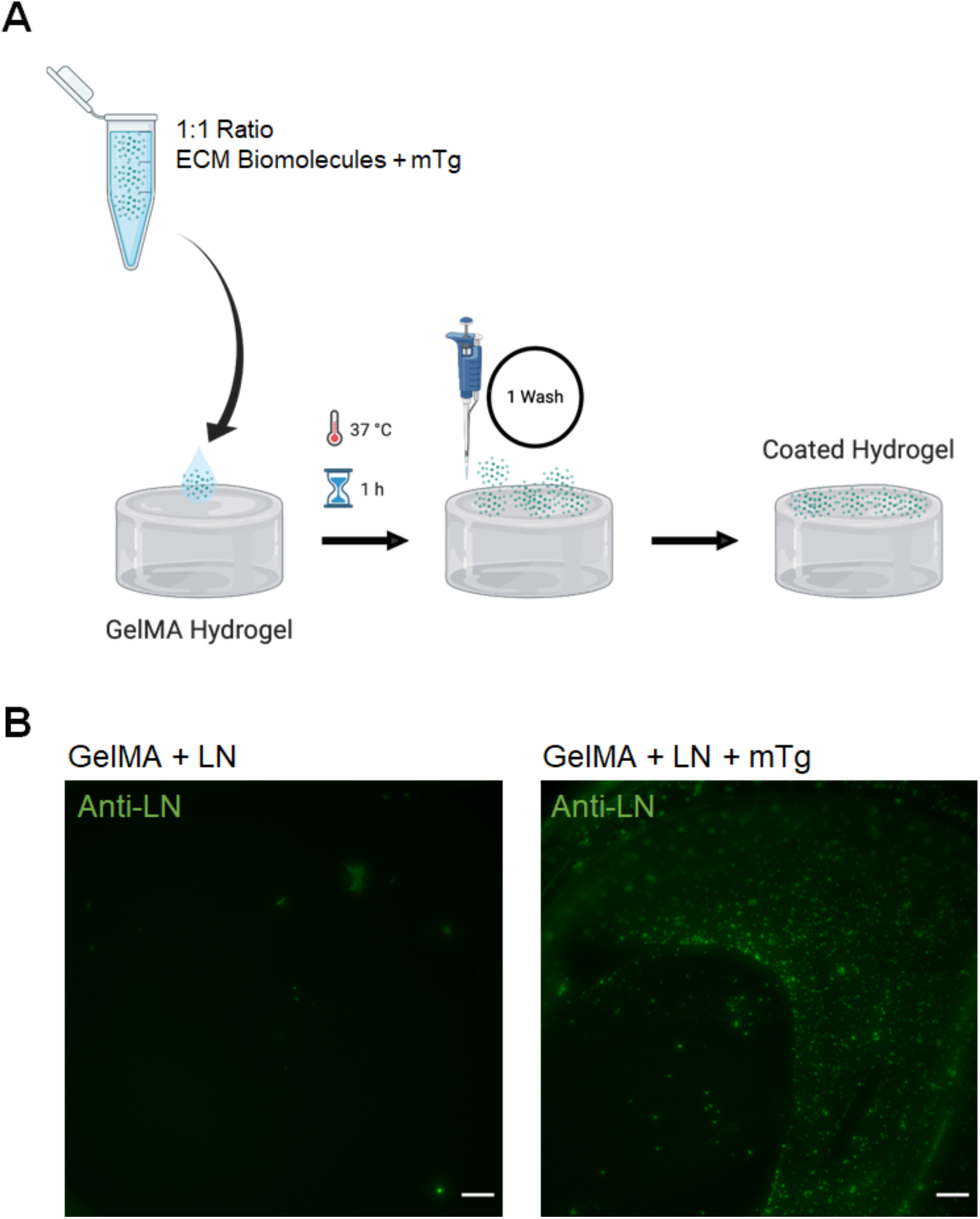
Methacrylamide-functionalized gelatin (GelMA) hydrogels are coated with extracellular matrix (ECM) biomolecules using microbial transglutaminase (mTg). **(A)** Experimental procedure for coating hydrogels. ECM biomolecules (10 μg/mL) and mTg (0.5 mg/mL) were mixed in a 1:1 ratio and pipetted onto GelMA hydrogels to coat the hydrogel surface. Created with BioRender.com. **(B)** GelMA hydrogels coated with laminin (LN) by adsorption or using the mTg protocol demonstrate significantly increased protein attachment using mTg. Green: laminin. Scale bar: 200 μm.

### 3.3. Epithelial cells cultured on ECM biomolecule combinations show variation in cell attachment and CK18 intensity

We subsequently assessed EEC attachment and CK18 expression on GelMA hydrogels coated with mTg-immobilized ECM proteins and proteoglycans to evaluate whether combinations down-selected from the microarray experiments would facilitate improved EEC attachment or CK18 expression. EEC attachment and CK18 expression were evaluated relative to control GelMA hydrogels as well as GelMA hydrogels coated with fibronectin or collagen IV + laminin factors previously used in the literature as endometrial basement membrane mimics [3]. We first examined cell attachment at CK18 expression using hits with highest cell adhesion in 2D microarray experiments (C4+TC: Collagen IV + Tenascin C; C1+C3: Collagen 1 + Collagen 3). Although overall analysis of the number of adhered cells suggested significant differences between conditions (Welch’s ANOVA, *p*=0.018), post hoc analysis showed no differences between experimental (Collagen IV + Tenascin C; Collagen I + Collagen III) and controls (Fig. 3A). CK18 intensity was not found to differ between groups (Fig. 3B; One-way ANOVA, *p*=0.88). We then examined the role of ECM combinations that resulted in the highest CK18 intensity in 2D microarray experiments (HA+TC: Hyaluronic Acid + Tenascin C; C5: Collagen V), finding no differences in the number of cells attached (Fig. 4A; One-way ANOVA, *p*=0.77) or in CK18 intensity (Fig. 4B; One-way ANOVA, *p*=0.31) on functionalized GelMA hydrogels. Finally, comparing ECM combinations with median cell attachment and CK18 intensity in 2D microarray experiments (C1+HA: Collagen I + Hyaluronic Acid; C5+HA: Collagen V + Hyaluronic Acid), high cell attachment and median CK18 intensity (C1: Collagen I), or median cell attachment and high CK18 intensity (C3: Collagen III), we observed no significant differences in cell attachment (Fig. 5A; Welch’s ANOVA, *p*=0.17). However, CK18 intensity was statistically significantly different between groups (Fig. 5B; Welch’s ANOVA, *p*=4.8×10^−4^), with post hoc analysis revealing significantly increased CK18 intensity in collagen I versus fibronectin coatings.

**Figure 3.**
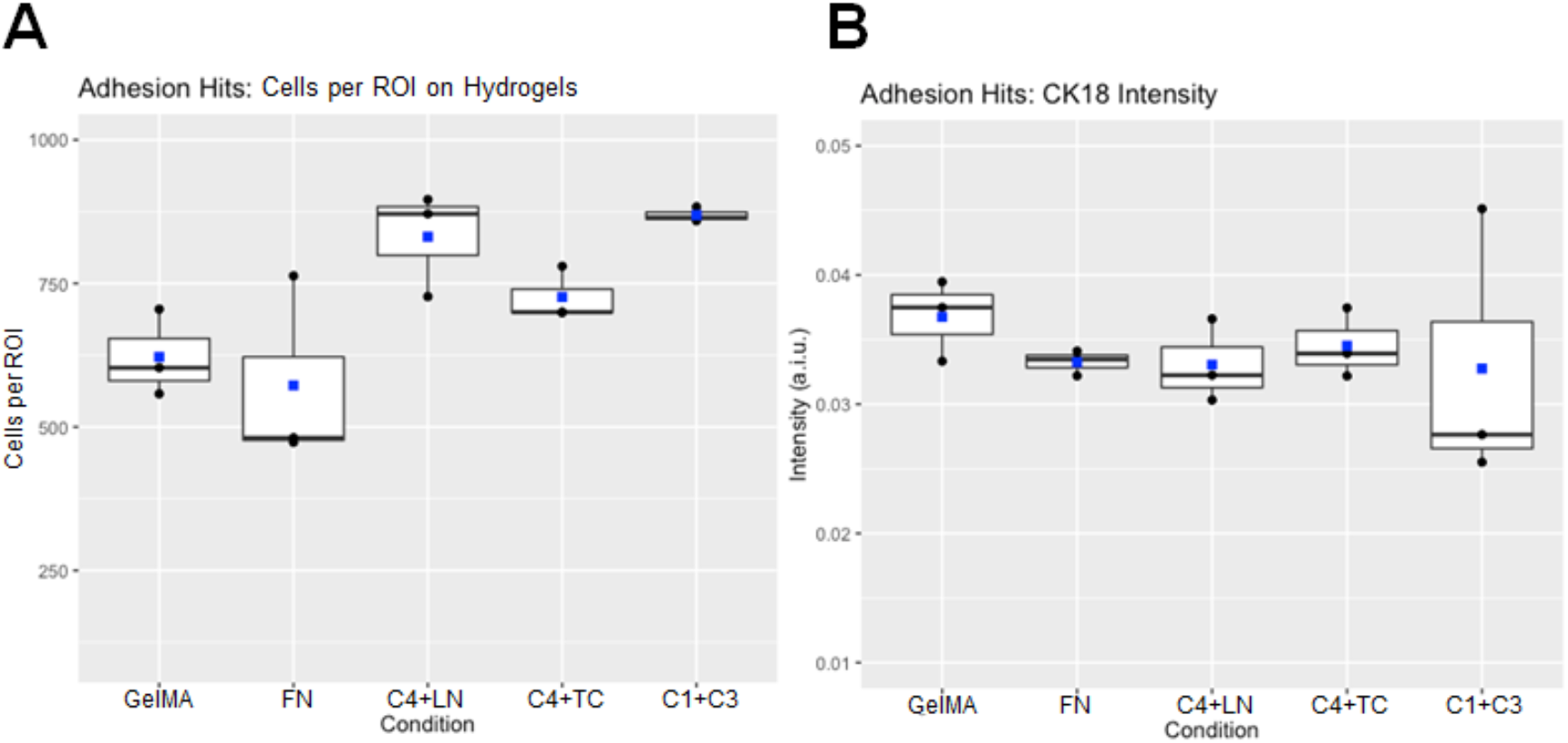
Cell attachment and cytokeratin 18 (CK18) intensity on methacrylamide-functionalized gelatin (GelMA) hydrogels coated with extracellular matrix combinations with highest adhesion in microarray experiments (C4+TC: Collagen IV + Tenascin C and C1+C3: Collagen 1 + Collagen 3). Control conditions consist of GelMA with no coating (GelMA) and coating conditions from literature (FN: Fibronectin and C4+LN: Collagen IV + Laminin). Data consists of n=3 hydrogels per condition (1 ROI per gel) with 1 maximum intensity confocal image analyzed per hydrogel. Data presented in box plots with blue squares representing the mean. **(A)** Average number of cells per ROI for each condition. Cells per ROI showed significant (Welch’s ANOVA: *p*=0.018) differences between groups. **(B)** Average CK18 intensity of cells on hydrogels for each condition. CK18 intensity showed no differences between groups (One-way ANOVA: *p*=0.88).

**Figure 4.**
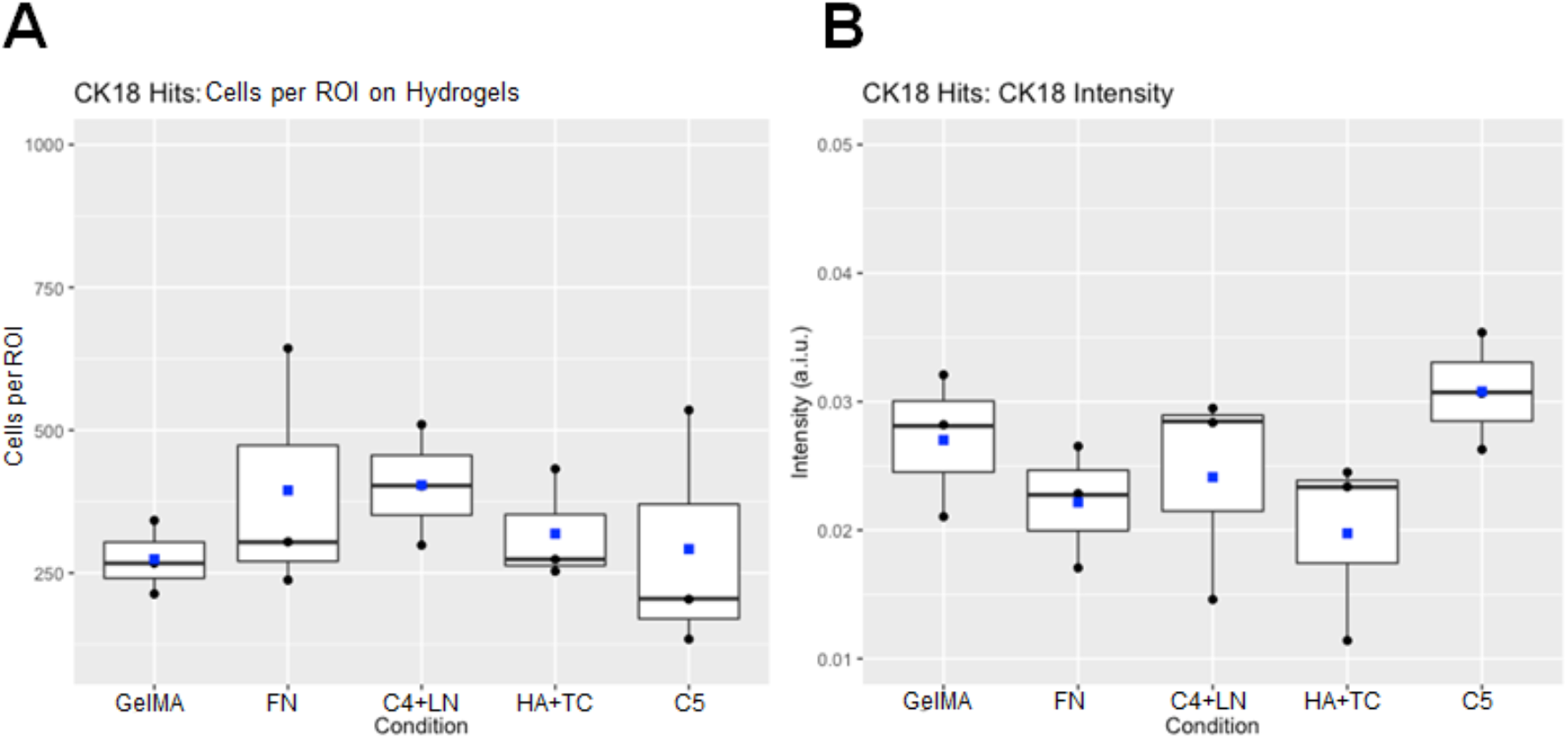
Cell attachment and cytokeratin 18 (CK18) intensity on methacrylamide-functionalized gelatin (GelMA) hydrogels coated with extracellular matrix combinations with highest CK18 intensity in microarray experiments (HA+TC: Hyaluronic Acid + Tenascin C and C5: Collagen 5). Control conditions consist of GelMA with no coating (GelMA) and coating conditions from literature (FN: Fibronectin and C4+LN: Collagen IV + Laminin). Data consists of n=3 hydrogels per condition (1 ROI per gel) with 1 maximum intensity confocal image analyzed per hydrogel. Data presented in box plots with blue squares representing the mean. **(A)** Average number of cells per ROI for each condition. Cells per ROI showed no differences between groups (Welch’s ANOVA: *p*=0.77). **(B)** Average CK18 intensity of cells on hydrogels for each condition. CK18 intensity showed no differences between groups (One-way ANOVA: *p*=0.31).

**Figure 5.**
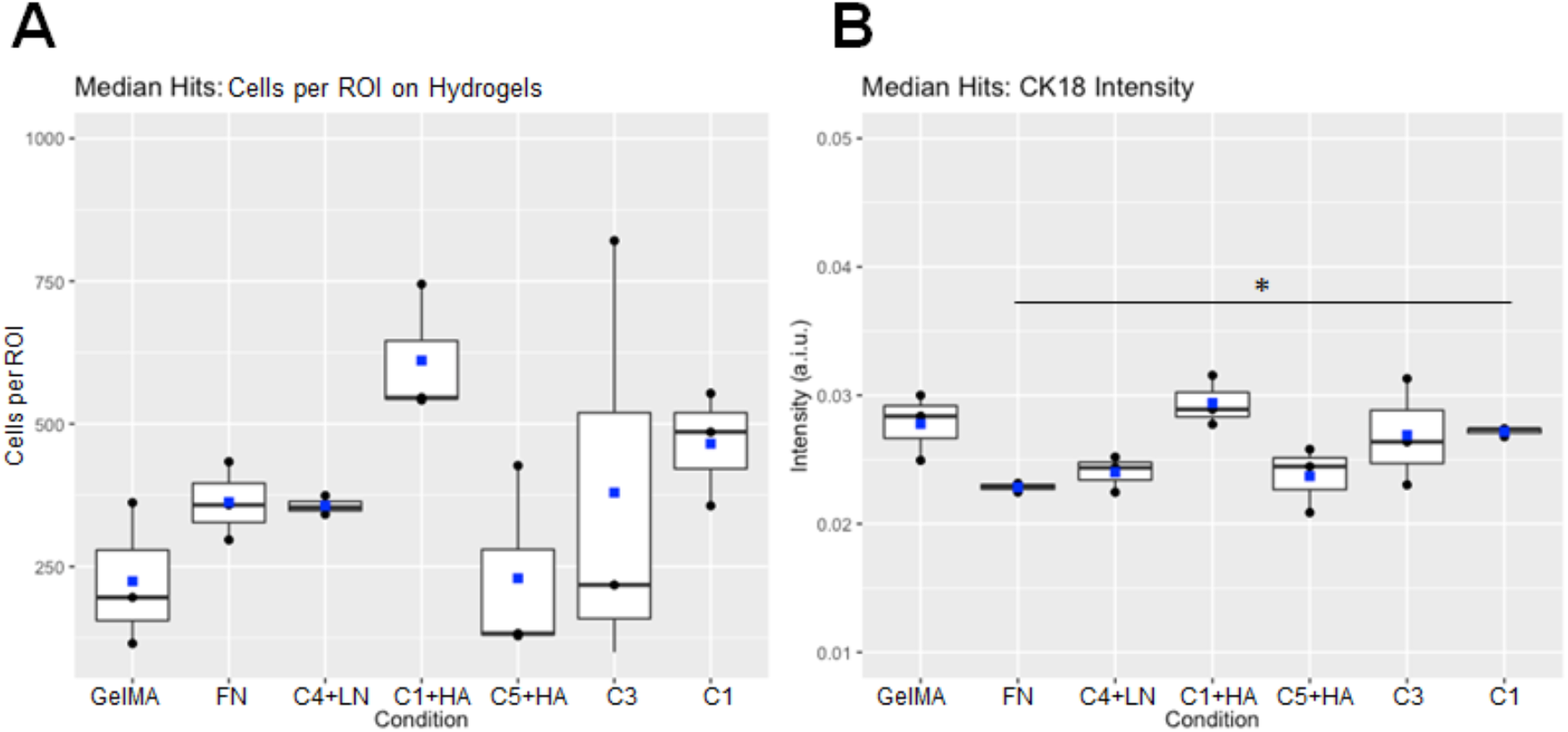
Cell attachment and cytokeratin 18 (CK18) intensity on methacrylamide-functionalized gelatin (GelMA) hydrogels coated with extracellular matrix combinations with median adhesion and CK18 intensity (C1+HA: Collagen I + Hyaluronic Acid and C5+HA: Collagen V + Hyaluronic Acid), high cell adhesion and median CK18 intensity (C1: Collagen I), and median cell adhesion and high CK18 intensity (C3: Collagen III) in microarray experiments. Control conditions consist of GelMA with no coating (GelMA) and coating conditions from literature (FN: Fibronectin and C4+LN: Collagen IV + Laminin). Data consists of n=3 hydrogels per condition (1 ROI per gel) with 1 maximum intensity confocal image analyzed per hydrogel. Data presented in box plots with blue squares representing the mean. **(A)** Average number of cells per ROI for each condition. Cells per ROI showed no differences between groups (Welch’s ANOVA: *p*=0.17). **(B)** Average CK18 intensity of cells on hydrogels for each condition. CK18 intensity showed statistically significant differences between groups (Welch’s ANOVA: *p*=4.8×10^−4^) with post hoc analysis demonstrating that CK18 intensity was significantly increased for collagen I vs fibronectin conditions.

### 3.4. Epithelial cells deposit nascent proteins onto GelMA hydrogels

We evaluated the potential for EECS to significantly remodel their basement membrane environment on 3D GelMA hydrogels, examining nascent protein deposition in response to ECM combinations that increased cell attachment in microarray experiments (C4+TC: Collagen IV + Tenascin C; C1+C3: Collagen I + Collagen III). Within 1 hour, EECs deposited observable levels of nascent proteins onto hydrogels in all conditions (Fig. 6A), with analysis of the intensity of nascent protein deposition revealed no difference between groups (Fig. 6B; One-way ANOVA, *p*=0.24).

**Figure 6.**
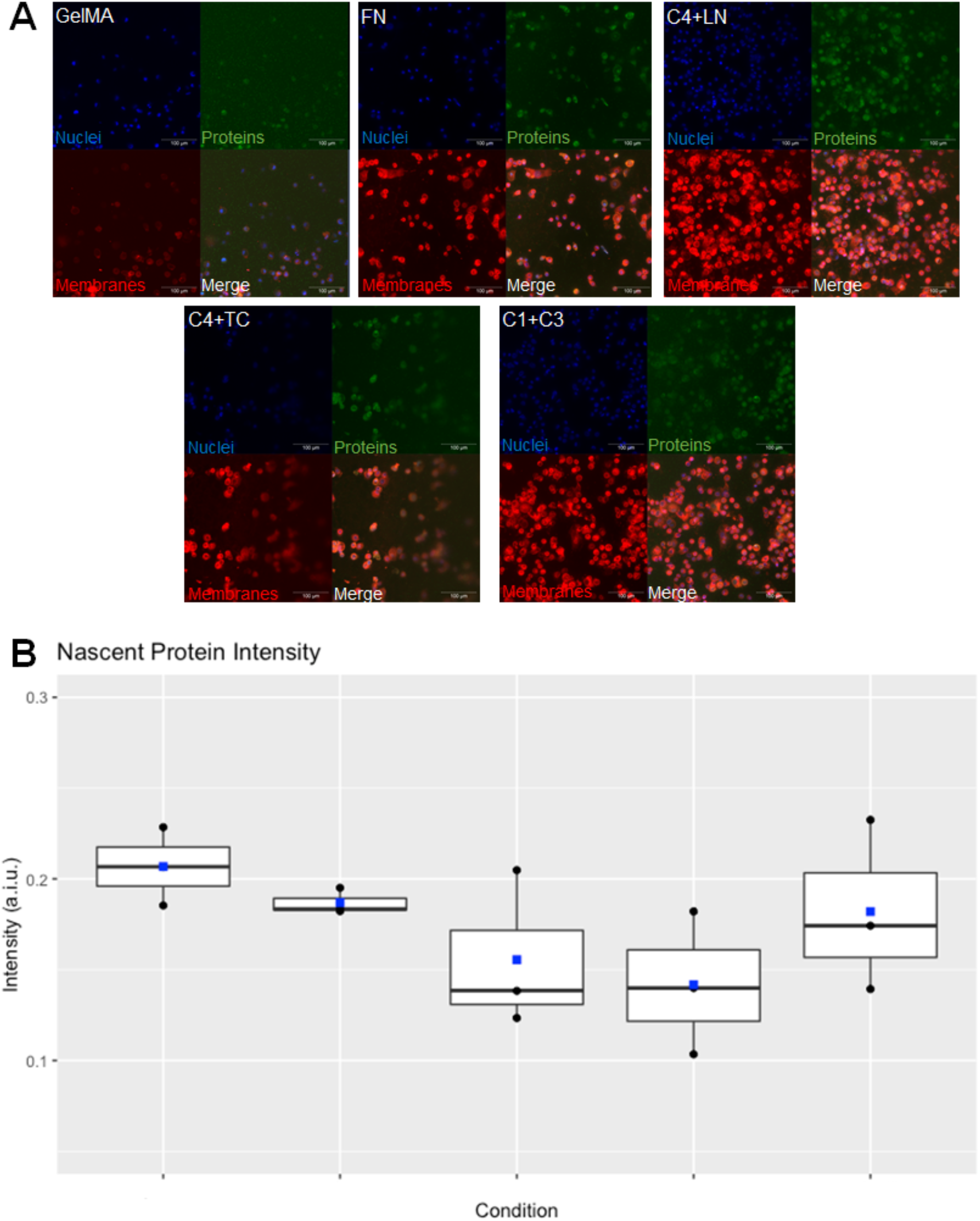
Primary endometrial epithelial cells deposit nascent proteins onto methacrylamide-functionalized gelatin (GelMA) hydrogels. **(A)** Nascent protein deposition after 1 hour on extracellular matrix (ECM) combinations highest adhesion in microarray experiments (C4+TC: Collagen IV + Tenascin C and C1+C3: Collagen 1 + Collagen 3). Control conditions consist of GelMA with no coating (GelMA) and coating conditions from literature (FN: Fibronectin and C4+LN: Collagen IV + Laminin). Green: nascent proteins; Red: cell membranes; Blue: nuclei. Scale: 100 μm. **(B)** Nascent protein intensity for each condition (no significant differences; One-way ANOVA: *p*=0.24). Data consists of n=3 hydrogels per condition (1 ROI per gel) with 1 maximum intensity confocal image analyzed per hydrogel. Data presented in box plots with blue squares representing the mean.

## 4. Discussion

Dynamic changes in the endometrial ECM plays a key role in homeostasis, pregnancy, and endometrial-associated pathologies [1]. Previous studies have identified a wide range of ECM-associated proteins and proteoglycans in the endometrium and decidua whose expression changes markedly across the menstrual cycle or in response to steroidal sex hormones [1]. Nevertheless, developing a deeper understanding of the role of the endometrial basement membrane and extracellular matrix on endometrial epithelial cell behavior may provide critical information on cell attachment phenotype (e.g., cytokeratin expression). Understanding endometrial epithelial cell attachment behavior to the basement membrane would also provide crucial information on how trophectoderm signaling and alterations to the basement membrane may facilitate blastocyst adhesion, disruption, breach, and resealing of the endometrial epithelium. Our goal was to identify ECM combinations that facilitate improved endometrial epithelial cell attachment and CK18 expression in order to support the development of tissue engineering systems that more appropriately mimic the endometrial ECM.

We have previously described overlaid cultures of endometrial epithelial cells on GelMA hydrogels [25]; however, we observed instability of the epithelial cultures over time and inability of the cultures to form a confluent monolayer. We hypothesized that an appropriate basement membrane layer on the surface of the GelMA hydrogel may be required to form a consistent, confluent monolayer. To determine whether a broader set of ECM biomolecule combinations inspired by the composition of the native endometrium may impact the stability of the epithelial layers, we performed high throughput experiments using microarrays to assess the effect of 55 single and pairwise combinations of 10 ECM biomolecules found in the endometrium (collagens I, III, IV, V, decorin, fibronectin, hyaluronic acid, laminin, lumican, and tenascin C) on endometrial epithelial cell attachment and CK18 expression.

We found collagen IV + tenascin C and collagen I + collagen III combinations resulted in the highest number of EECs attached while hyaluronic acid + tenascin C and collagen V had the highest values for CK18 intensity. Further, collagen I + hyaluronic acid and collagen V + hyaluronic acid had values for cell attachment and CK18 intensity around the 15^th^ position from the highest values, while collagen I promoted high cell attachment and median CK18 intensity, while collagen III promoted median cell attachment and high CK18 intensity. These results identified combinations of key ECM proteins and proteoglycans found in the endometrium that affect EEC attachment and CK18 expression. For example, collagens I, III, and V are all present in the endometrium but collagen I is most prevalent during the proliferative and secretory phases, collagen III is present during all phases, and collagen V increases during decidualization [1]. Tenascin C is present near stromal cells surrounding proliferating or developing endometrial epithelia [1]. Logically, tenascin C would be relevant to these studies because EECs would likely be proliferating and developing in these cultures. Finally, hyaluronic acid may influence hydration of the tissue during the mid-proliferative and mid-secretory phases [1]. Interestingly, our microarray data did not identify fibronectin or collagen IV + laminin as combinations that resulted in high cell attachment or CK18 expression, despite many studies utilizing these as basement membrane mimics based on their prevalence in the endometrial basement membrane [1, 3]. Future efforts looking at a larger screen of metrics of EEC bioactivity may be required to understand the role of these biomolecules on endometrial epithelial cell activity.

We used these data to identify a set of biomolecules to immobilize on the surface of three-dimensional GelMA hydrogels to assess epithelial cell attachment and CK18 expression. We adapted a recently reported microbial transglutaminase (mTg) biomolecule immobilization strategy [16] to covalently bind biomolecules to GelMA hydrogels. This approach enables attachment of ECM combinations using a relatively simple protocol with immobilized biomolecules showing extended stability (28 days) in culture [16]. We then quantified cell attachment and CK18 intensity on GelMA hydrogels coated with a range of endometrial-inspired matrix biomolecules. While the number of cells per island and CK18 expression levels varied for each set of selected ECM biomolecules, the statistical significance of the data was variable.

Compared to controls (GelMA only, Fibronectin, Collagen IV + Laminin), matrix hits for adhesion Collagen IV + Tenascin C and Collagen I + Collagen III) had statistically different number of cells per island but not CK18 intensity. For combinations resulting in the highest CK18 intensity in microarray experiments (Hyaluronic Acid + Tenascin C and Collagen V), the numbers of cells attached and the CK18 intensity were not statistically significantly different between groups. Finally, for ECM combinations around the 15^th^ position from the highest values of cell attachment and CK18 intensity (Collagen I + Hyaluronic Acid and Collagen V + Hyaluronic Acid), high cell attachment and median CK18 intensity (Collagen I), and median cell attachment and high CK18 intensity (Collagen III), we determined that the cells attached did not statistically differ between groups but, CK18 intensity was statistically significantly different between groups. Post hoc analysis revealed statistical significance between CK18 intensity between the fibronectin and collagen I conditions. These data demonstrated that there is variation in cell response based on ECM coating and that CK18 expression, a marker of endometrial cell phenotype, was more strongly affected that overall numbers of attached cells. Taken together, these data suggest that careful selection of basement membrane ECM combinations must be considered for tissue engineering constructs under development to replicate the stratified endometrium.

Finally, we demonstrated functional metrics of EEC-mediated remodeling of the engineered basement membrane environment by quantifying nascent protein deposition by EECs on GelMA hydrogels. We observed rapid (~1 h) protein deposition, demonstrating EECs rapidly deposit their own ECM onto the surface on which they are cultured and continue to deposit proteins throughout their culture period regardless of the ECM biomolecules on which they are growing. These results suggest significant opportunities to examine variations in the specific proteins being deposited as well as shifts in matrix deposition as a function of initial basement membrane content. Extending on our previous use of the GelMA hydrogel platform to evaluate endometrial stromal models [14], these findings suggest opportunities to evaluate crosstalk between endometrial epithelial layers and underlying endometrial perivascular models as well as the opportunity to evaluate the role of external stimuli such as steroidal sex hormones in a fully stratified endometrial culture platform. We also recognize some limitations and future opportunities for these studies. The cells used in these studies are primary endometrial epithelial cells derived from a single donor. What matrix the EECs prefer may have donor-to-donor variability and may depend on the menstrual cycle phase during which the cells were collected as well as donor-to-donor cell preferences. For example, *Cook et al*. previously identified differences in epithelial cell behavior that was dependent on cell donors [3]. These results suggest that the basement membrane and ECM biomolecule combinations may vary between individuals and underlie the need for an adaptable tissue engineering approach such as the 2D microarray to 3D biomaterial pipeline described here, to help identify how this variability affects uterine function and health. Future studies that incorporate cells from multiple donors collected at various points in the menstrual cycle may provide insight in donor-to-donor and cycle-dependent variability in cell response to different ECM components.

## 5. Conclusions

The endometrium is a highly dynamic tissue that suggests the need for a dynamic model system to replicate processes of growth, remodeling, and breakdown in order to properly recapitulate endometrial physiology. The ability to create an endometrial basement membrane mimic within a tissue engineered construct would provide opportunities to develop complex platforms that mimic not just a single menstrual cycle phase but also various points in the menstrual cycle. Here, we demonstrate differential response of EECs to ECM biomolecule combinations using a coordinated set of high-throughput two-dimensional microarrays and three-dimensional matrix-functionalized GelMA hydrogels. We report an approach to coat GelMA hydrogels with combinations of ECM proteins and proteoglycan to investigate the role of engineered basement membrane composition on endometrial epithelial cell attachment and phenotypic markers via immunostaining as well as evaluate endometrial epithelial cell mediated matrix remodeling via nascent protein deposition. Together, these results suggest an approach to replicate features of a stratified endometrial model, notably epithelial cell adhesion and remodeling via combinations of immobilized ECM biomolecules that can be tuned to match the changing endometrial microenvironment.

## Acknowledgements

Research reported was supported by the National Institutes of Diabetes and Digestive and Kidney Diseases of the National Institutes of Health under Award Numbers R01 DK0099528 (B.A.C.H.) and R01 DK125471 (G.H.U.), the National Cancer Institute of the National Institutes of Health under Award Number R01 CA256481 (B.A.C.H), and by the National Institute of Biomedical Imaging and Bioengineering of the National Institutes of Health under Award Number T32 EB019944 (S.G.Z.). The content herein is solely the responsibility of the authors and does not necessarily represent the official views of the National Institutes of Health. The authors also gratefully acknowledge additional funding provided by the Department of Chemical & Biomolecular Engineering and the Carl R. Woese Institute for Genomic Biology at the University of Illinois at Urbana-Champaign. The authors also thank the Institute for Genomic Biology Core Facilities (Dr. Austin Cyphersmith) at the University of Illinois Urbana-Champaign for assistance with confocal imaging.

## Author Contributions

We describe contributions to the manuscript using the Contributor Roles Taxonomy (CRediT) [30, 31]: *Writing – Original Draft*: SGZ and IJ; *Writing – Review & Editing:* SGZ, IJ, KBHC, GHU, and BACH; *Conceptualization:* SGZ, KBHC, and BACH; *Investigation:* SGZ and IJ; *Methodology:* SGZ and IJ; *Formal Analysis:* SGZ and IJ; *Data Curation:* SGZ and IJ; *Visualization:* SGZ and IJ; *Project Administration:* BACH; *Resources:* KBHC, GHU, and BACH; *Funding Acquisition*: GHU and BACH; Supervision: KBHC, GHU, and BACH.

## Data Availability

The raw data required to reproduce these findings are available per request by contacting the corresponding author. The processed data required to reproduce these findings are available per request by contacting the corresponding author.

## References

[1] J.D. Aplin, A.T. Fazleabas, S.R. Glasser, L.C. Giudice, The Endometrium, Second ed., Informa Healthcare, United Kingdom, 2008.

[2] H. Singh, J.D. Aplin, Adhesion molecules in endometrial epithelium: tissue integrity and embryo implantation, J Anat 215(1) (2009) 3–13.

[3] C.D. Cook, A.S. Hill, M. Guo, L. Stockdale, J.P. Papps, K.B. Isaacson, D.A. Lauffenburger, L.G. Griffith, Local remodeling of synthetic extracellular matrix microenvironments by co-cultured endometrial epithelial and stromal cells enables long-term dynamic physiological function, Integr Biol (Camb) 9(4) (2017) 271–289.

[4] H. Wang, F. Pilla, S. Anderson, S. Martinez-Escribano, I. Herrer, J.M. Moreno-Moya, S. Musti, S. Bocca, S. Oehninger, J.A. Horcajadas, A novel model of human implantation: 3D endometrium-like culture system to study attachment of human trophoblast (Jar) cell spheroids, Mol Hum Reprod 18(1) (2012) 33–43.

[5] H. Wang, S. Bocca, S. Anderson, L. Yu, B.S. Rhavi, J. Horcajadas, S. Oehninger, Sex steroids regulate epithelial-stromal cell cross talk and trophoblast attachment invasion in a three-dimensional human endometrial culture system, Tissue Eng Part C Methods 19(9) (2013) 676–87.

[6] C.M. Oefner, A. Sharkey, L. Gardner, H. Critchley, M. Oyen, A. Moffett, Collagen type IV at the fetal-maternal interface, Placenta 36(1) (2015) 59–68.

[7] P.D. Yurchenco, Basement membranes: cell scaffoldings and signaling platforms, Cold Spring Harb Perspect Biol 3(2) (2011).

[8] R. Cruz-Acuna, A.J. Garcia, Synthetic hydrogels mimicking basement membrane matrices to promote cell-matrix interactions, Matrix Biol 57-58 (2017) 324–333.

[9] A.M. Carter, Animal models of human placentation--a review, Placenta 28 Suppl A (2007) S41–7.

[10] A.M. Carter, A.M. Mess, Mammalian Placentation: Implications for Animal Models, Pathobiology of Human Disease 2014, pp. 2423–2442.

[11] J. Cha, X. Sun, S.K. Dey, Mechanisms of implantation: strategies for successful pregnancy, Nat Med 18(12) (2012) 1754–67.

[12] C.S. Hughes, L.M. Postovit, G.A. Lajoie, Matrigel: a complex protein mixture required for optimal growth of cell culture, Proteomics 10(9) (2010) 1886–90.

[13] E.A. Aisenbrey, W.L. Murphy, Synthetic alternatives to Matrigel, Nature Reviews Materials 5(7) (2020) 539–551.

[14] S.G. Zambuto, K.B.H. Clancy, B.A.C. Harley, A gelatin hydrogel to study endometrial angiogenesis and trophoblast invasion, Interface Focus 9(5) (2019).

[15] S. Pedron, B.A. Harley, Impact of the biophysical features of a 3D gelatin microenvironment on glioblastoma malignancy, J Biomed Mater Res A 101(12) (2013) 3404–15.

[16] R.R. Besser, A.C. Bowles, A. Alassaf, D. Carbonero, I. Claure, E. Jones, J. Reda, L. Wubker, W. Batchelor, N. Ziebarth, R. Silvera, A. Khan, R. Maciel, M. Saporta, A. Agarwal, Enzymatically crosslinked gelatin-laminin hydrogels for applications in neuromuscular tissue engineering, Biomater Sci 8(2) (2020) 591–606.

[17] Y. Aratyn-Schaus, P.W. Oakes, J. Stricker, S.P. Winter, M.L. Gardel, Preparation of Complaint Matrices for Quantifying Cellular Contraction, JoVE (Journal of Visualized Experiments) 46 (2010).

[18] J.R. Tse, A.J. Engler, Preparation of hydrogel substrates with tunable mechanical properties, Current protocols in cell biology 47(1) (2010) 10–16.

[19] J.H. Wen, L.G. Vincent, A. Fuhrmann, Y.S. Choi, K.C. Hribar, H. Taylor-Weiner, S. Chen, A.J. Engler, Interplay of matrix stiffness and protein tethering in stem cell differentiation, Nature Materials 13(10) (2014) 979–987.

[20] C.J. Flaim, J.S. Chien, S.N. Bhatia, An extracellular matrix microarray for probing cellular differentiation, Nature Methods 2(2) (2005) 119–125.

[21] D.A. Brafman, S. Chien, K. Willert, Arrayed cellular microenvironments for identifying culture and differentiation conditions for stem, primary and rare cell populations, Nature Protocols 7(4) (2012) 703–717.

[22] K.B. Kaylan, V. Ermilova, R.C. Yada, G.H. Underhill, Combinatorial microenvironmental regulation of liver progenitor differentiation by Notch ligands, TGFβ and extracellular matrix, Scientific Reports 6(1) (2016) 1–15.

[23] J. Schindelin, I. Arganda-Carreras, E. Frise, V. Kaynig, M. Longair, T. Pietzsch, S. Preibisch, C. Rueden, S. Saalfeld, B. Schmid, J.Y. Tinevez, D.J. White, V. Hartenstein, K. Eliceiri, P. Tomancak, A. Cardona, Fiji: an open-source platform for biological-image analysis, Nat Methods 9(7) (2012) 676–82.

[24] C. McQuin, A. Goodman, V. Chernyshev, L. Kamentsky, B.A. Cimini, K.W. Karhohs, M. Doan, L. Ding, S.M. Rafelski, D. Thirstrup, W. Wiegraebe, S. Singh, T. Becker, J.C. Caicedo, A.E. Carpenter, CellProfiler 3.0: Next-generation image processing for biology, PLoS Biol 16(7) (2018).

[25] S.G. Zambuto, K.B.H. Clancy, B.A.C. Harley, Tuning Trophoblast Motility in a Gelatin Hydrogel via Soluble Cues from the Maternal-Fetal Interface, Tissue Eng Part A (2020).

[26] M. De Colli, M. Massimi, A. Barbetta, B.L. Di Rosario, S. Nardecchia, L. Conti Devirgiliis, M. Dentini, A biomimetic porous hydrogel of gelatin and glycosaminoglycans cross-linked with transglutaminase and its application in the culture of hepatocytes, Biomed Mater 7(5) (2012) 055005.

[27] G. Damodaran, R. Collighan, M. Griffin, A. Pandit, Tethering a laminin peptide to a crosslinked collagen scaffold for biofunctionality, J Biomed Mater Res A 89(4) (2009) 1001–10.

[28] C. Loebel, R.L. Mauck, J.A. Burdick, Local nascent protein deposition and remodelling guide mesenchymal stromal cell mechanosensing and fate in three-dimensional hydrogels, Nat Mater 18(8) (2019) 883–891.

[29] C. Loebel, M.Y. Kwon, C. Wang, L. Han, R.L. Mauck, J.A. Burdick, Metabolic Labeling to Probe the Spatiotemporal Accumulation of Matrix at the Chondrocyte-Hydrogel Interface, Advanced Functional Materials (2020).

[30] Brand, L. Allen, M. Altman, M. Hlava, J. Scott, Beyond authorship: attribution, contribution, collaboration, and credit, Learned Publishing 28(2) (2015) 151–155.

[31] L. Allen, J. Scott, A. Brand, M. Hlava, M. Altman, Publishing: Credit where credit is due, Nature 508(7496) (2014) 312–3.

